# Identification of piRNA binding sites reveals the Argonaute regulatory landscape of the *C. elegans* germline

**DOI:** 10.1101/262113

**Authors:** En-Zhi Shen, Hao Chen, Ahmet R. Ozturk, Shikui Tu, Masaki Shirayama, Wen Tang, Yue-He Ding, Si-Yuan Dai, Zhiping Weng, Craig C. Mello

**Affiliations:** RNA Therapeutics Institute; Program in Bioinformatics and Integrative Biology University of Massachusetts Medical School, Worcester, MA 01605, USA; Howard Hughes Medical Institute; Co-first author

## Abstract

piRNAs (Piwi-interacting small RNAs) engage Piwi Argonautes to silence transposons and promote fertility in animal germlines. Genetic and computational studies have suggested that *C. elegans* piRNAs tolerate mismatched pairing and in principle could target every transcript. Here we employ *in vivo* cross-linking to identify transcriptome-wide interactions between piRNAs and target RNAs. We show that piRNAs engage all germline mRNAs and that piRNA binding follows microRNA-like pairing rules. Targeting correlates better with binding energy than with piRNA abundance, suggesting that piRNA concentration does not limit targeting. In mRNAs silenced by piRNAs, secondary small RNAs accumulate at the center and ends of piRNA binding sites. In germline-expressed mRNAs, however, targeting by the CSR-1 Argonaute correlates with reduced piRNA binding density and suppression of piRNA-associated secondary small RNAs. Our findings reveal physiologically important and nuanced regulation of individual piRNA targets and provide evidence for a comprehensive post transcriptional regulatory step in germline gene expression.

## INTRODUCTION

Argonaute (AGO) proteins and their associated small RNAs are fundamental regulators of transcriptional and post-transcriptional gene regulation(Czech and Hannon, 2011; Ghildiyal and Zamore, 2009; Hutvagner and Simard, 2008; Meister, 2013; Siomi and Siomi, 2009; Thomson and Lin, 2009). Piwi proteins are members of the RNaseH-related Argonaute superfamily and associate with small RNAs (i.e., Piwi-interacting RNAs or piRNAs) to form piRNA-induced silencing complexes (piRISCs)(Czech and Hannon, 2016; Iwasaki et al., 2015; Juliano et al., 2011; Malone and Hannon, 2009; Weick and Miska, 2014). The genomic origins, sequences, and lengths of animal piRNAs vary, but some of the biological functions of piRISCs appear to be shared. For example, piRISCs are required for fertility and transposon silencing in worms, flies, and mice(Aravin et al., 2001, 2007; Batista et al., 2008; Carmell et al., 2007; Cox et al., 1998; Houwing et al., 2007; Lee et al., 2012; Lin and Spradling, 1997; Savitsky et al., 2006; Siomi et al., 2011; Thomson and Lin, 2009; Vagin et al., 2006). A growing number of studies suggest that piRNAs and Piwi Argonautes may regulate many, if not all, germline mRNAs (Fagegaltier et al., 2016; Gou et al., 2014; Vourekas et al., 2016; Zhang et al., 2015b).

The *C. elegans* Piwi protein, PRG-1, binds an abundant class of germline-expressed 21-nucleotide (nt) piRNAs with a 5’ uridine (21U-RNAs)(Batista et al., 2008; Ruby et al., 2006). Targeting by the 21U-RNA/PRG-1 piRISC complex recruits an RNA-dependent RNA polymerase (RdRP) that initiates the *de novo* synthesis of secondary 22-nt small RNAs that are templated directly from the target RNA and exhibit a bias for a 5’ guanosine residue. These so-called 22G-RNAs engage an expanded group of <u>w</u>orm <u>A</u>r<u>go</u>nautes (WAGOs) that function downstream of piRNAs to silence transposons and many endogenous genes. The WAGO pathway is required for long-term maintenance of silencing(Bagijn et al., 2012; Lee et al., 2012).

piRNA targeting in *C. elegans* permits mismatches, suggesting that thousands of endogenous mRNAs could be targeted by piRNAs (Lee et al., 2012). However, anti-silencing mechanisms are thought to prevent or reduce the sensitivity of endogenous mRNAs to piRNA-mediated silencing. The CSR-1 pathway, for example, is thought to be one arm of a “self” recognition pathway that protects endogenous mRNAs from piRNA surveillance (Seth et al., 2013). CSR-1 engages RdRP-derived small RNAs templated from nearly all germline-expressed mRNAs (Claycomb et al., 2009). However, it is unknown whether CSR-1 blocks PRG-1 targeting directly or if it acts downstream to prevent WAGO recruitment.

The identification of piRNA targets is essential for deciphering the roles of piRNAs in both sequence-directed immunity and more broadly in the regulation of germline gene expression. Here, we optimize a crosslinking, ligation, and sequencing of hybrids (CLASH) protocol to identify piRNAs and associated (candidate) target RNA binding sites in *C. elegans* (Helwak et al., 2013; Van Nostrand et al., 2016; Vourekas and Mourelatos, 2014). We identified 200,000 high-confidence piRNA–target site interactions. The overwhelming majority of interactions were between piRNAs and mRNAs. Bioinformatics analysis of the hybrids revealed that targets are enriched for energetically favorable Watson-Crick pairing with their associated piRNAs. We show that the seed sequence (i.e., positions 2 to 8) and supplemental nucleotides near the 3’ end (positions 14 to 19) of the piRNA are important determinants of piRNA-target binding and silencing, suggesting that piRNA targeting resembles miRNA targeting.

piRNA target sites defined by CLASH show a non-random pattern of WAGO 22G-RNAs that initiate at both ends and near the center (position 12) of the piRNA target site, consistent with local recruitment of RdRP. Analysis of CLASH hybrids obtained from CSR-1-depleted animals suggest that CSR-1 protects its targets from PRG-1 binding and WAGO-dependent silencing. Our findings reveal that the entire germline mRNA transcriptome engages piRISC, and suggests how germline Argonaute pathways are coordinated to achieve comprehensive regulation and surveillance of germline gene expression.

## RESULTS

### PRG-1 CLASH directly identifies piRNA-target chimeras

We used a modified Cross Linking and Selection of Hybrids (CLASH) approach to identify RNAs associated with the *C. elegans* PRG-1-piRISC complex. Briefly, CLASH involves the *in vivo* cross-linking of RNAs to a protein of interest followed by immunoprecipitation (IP), trimming of RNA ends, ligation to form hybrids between proximal RNAs within the crosslinked complex, cDNA preparation, library construction, and deep sequencing (see Experimental Procedures, Figure 1A-E). In principle, this procedure should allow the recovery of hybrid-sequence reads formed when piRNAs are ligated to proximal cellular target RNAs within the cross-linked PRG-1 IP complex.

**Figure 1.**
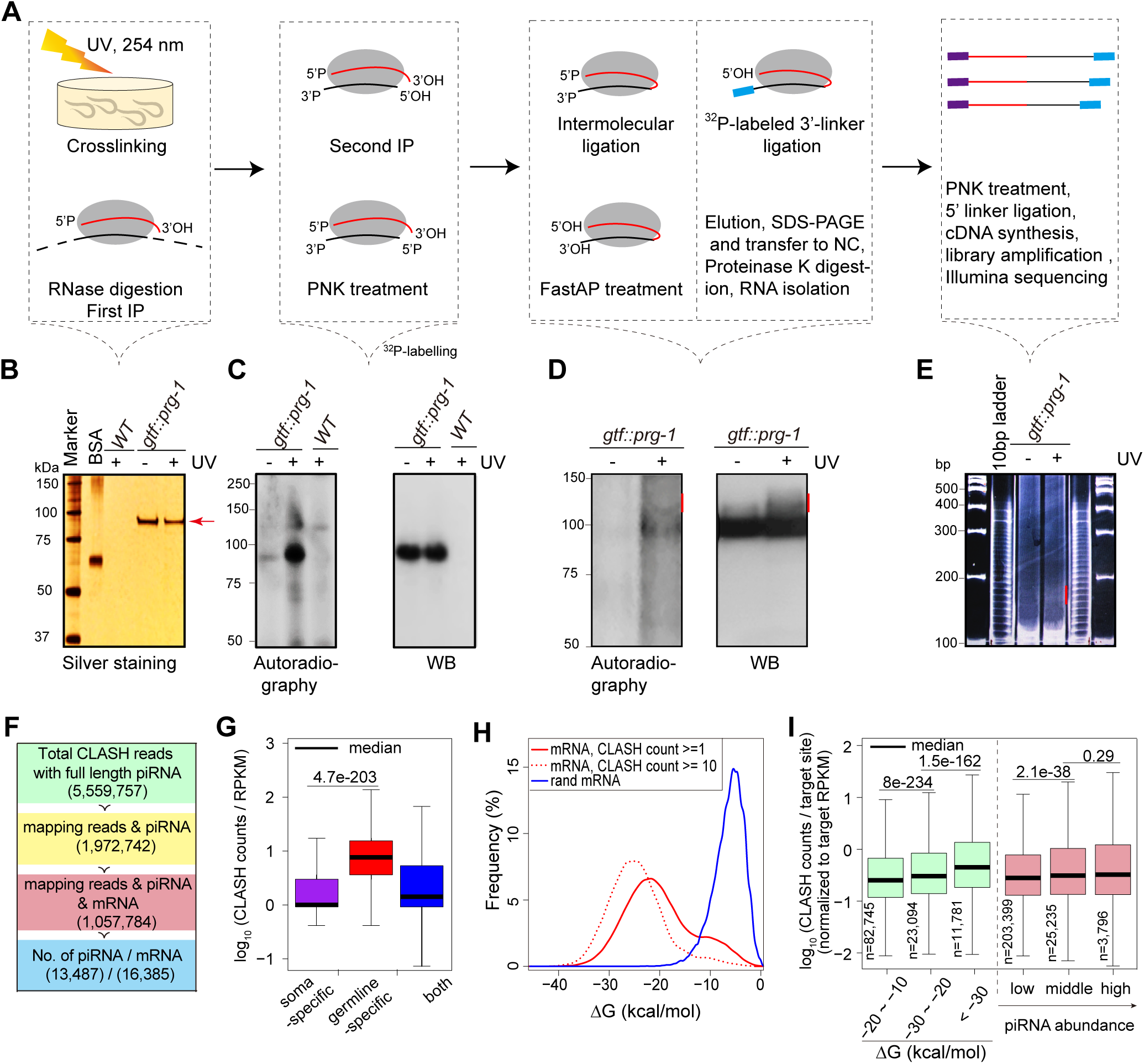
PRG-1 CLASH identifies piRNA–target chimeras in *C. elegans*. **(A)** CLASH workflow. See Experimental Procedures for details of steps. PRG-1 is indicated by gray oval, piRNA as red line, and mRNA as black line. Linkers are indicated as blue (3’) and purple (5’) rectangles. **(B)** Silver stain analysis of purified PRG-1 from wild type (WT) and *gfp::tev::flag::prg-1* worms with or without 254-nm UV irradiation. PRG-1 protein is indicated by red arrow. **(C-D)** Autoradiograph (left) and western blot (right) analyses of FLAG::PRG-1 complexes containing ^32^P-labeled RNA immunoprecipitated from lysates of worms with or without UV crosslinking. Panel D shows independent samples with red lines next to the region excised from the membrane. **(E)** Polyacrylamide gel showing the library amplification products (red line) isolated for Illumina sequencing. **(F)** Distribution of reads and mapping results from CLASH. **(G)** Normalized CLASH read counts per gene for soma-specific genes, germline-specific genes, and both. **(H)** Distribution of predicted binding energies between piRNAs and target sites identified by CLASH is stronger than between randomly matched pairs. **(I)** Box plots of the change in CLASH counts per target site for increasing piRNA:target base pairing (∆G, kcal/mol) and piRNA abundance (0–33%, 33–66%, 66–100%). Median is indicated by solid black lines. Each set is significantly different from the prior one, as indicated by the p-values. See also Figures S1

To perform CLASH, we first used CRISPR/Cas9 genome editing to introduce a GFP-TEV-FLAG (GTF) multiplex tag into the endogenous *prg-1* locus (Kim et al., 2014). In addition to direct fluorescence detection this tag also permits tandem affinity purification with a TEV protease-mediated elution after the first affinity step. GTF::PRG-1 exhibited a robust expression and was prominently localized in P-granules (Figure S1A) in a pattern identical to that previously reported in PRG-1 immunolocalization studies (Batista et al., 2008). Moreover, the GTF::PRG-1 fusion protein was functional, as evidenced by its ability to mediate piRNA-dependent silencing of a *gfp::cdk-1* reporter gene (Figure S1B).

We then carefully optimized each step of the CLASH procedure using GTF-PRG-1 (Figure 1 A-E; Experimental Procedures) (Broughton et al., 2016; Helwak et al., 2013). The tandem affinity purification of GTF::PRG-1 resulted in recovery of a single prominent protein of the expected size in silver-stained SDS-PAGE gels (Figure 1B). Although GTF::PRG-1 stably associated with piRNAs under these purification conditions, the recovery of longer associated RNA required the pretreatment of the worms with ultra-violet light. These crosslinked RNA Protein complexes, RNPs, were then treated with nuclease to trim the long RNAs, followed by intermolecular ligation to form RNA hybrids between the piRNA and target (Figure 1B-D). RNA hybrids of approximately 42 nts were recovered by gel purification (Figure 1 E) and were used for library construction and deep-sequencing.

In two independent experiments we found similar distributions of mapped sequence reads (Figure S1C-H). Together, these comprised a total of ~21million reads, including a total of ~7-million reads corresponding to 13,487 different piRNAs. Most of these piRNA-containing reads lacked a hybrid sequence (1,083,173), or the hybrid sequences could not be mapped to the genome because they were too short, or for other reasons (3,944,926). We obtained 2,106,796 hybrid reads composed of a piRNA sequence and a genome-mapping sequence, of which ~1.5 million were composed of a single piRNA sequence fused to an mRNA. In addition to mRNA chimeras, we detected piRNAs fused to sequences corresponding to rRNA (82,517 reads), tRNA (8,296 reads), pseudogenes (34,177 reads), lincRNA (1,845 reads), miRNA (1,334 reads), introns (9,006 reads), and transposable elements (14,468 reads).

### CLASH reveals piRNA target sites in germline mRNAs

Because mRNA chimeras were by far the most abundant type of hybrid read, we chose to focus on mRNA hybrids in the present study. Altogether a total of 16,385 genes were represented among the piRNA hybrids (Figure 1F). We found that “soma-specific” mRNAs were strongly under-represented in the CLASH data (Figure 1G) (Beanan and Strome, 1992; Li et al., 2014a), consistent with the idea that CLASH captures interactions between piRNAs and mRNAs that occur in the germline, and not interactions that occur in lysates. The frequency of recovery of each piRNA by CLASH correlated with its level in the input sample as measured by small RNA sequencing (Figure S1I, r = 0.58, P < 0.005).

The nuclease treatment during the CLASH procedure was optimized to produce chimeras of approximately 40 nucleotides. Thus, each chimera potentially reflects a piRNA/target mRNA duplex ligated at, or near, one end of the duplex. We noted, however, that not all chimeras contained a full-length piRNA and that the recovered target regions varied in length, indicating some variability in nuclease trimming during the CLASH procedure. Therefore, prior to searching for base-pairing interactions, we inferred the full-length piRNA and extended the empirically defined target space by adding nucleotides to each end, creating “ideal” piRNA/target RNA pairs (See Experimental Procedures).

We next predicted the most energetically favorable piRNA-mRNA interactions from in silico folding of these “ideal” sequences and compared it with predicted binding energies in a control data set (Figure 1H). This analysis showed that stable base-pair interactions were strongly enriched in the recovered piRNA-mRNAs chimeras. In fact, when normalized for mRNA levels, hybrid read counts per target site correlated better with binding energy than with piRNA abundance (see Discussion) (Figure 1I). Chimeras in which the piRNA 3’ end was contiguous with mRNA sequence were roughly 20-fold more frequent than chimeras ligated at piRNA 5’ ends (Figure S1J). These findings are consistent with the idea that, within piRISC, piRNA 3’ ends are more available to interact with mRNA fragments. Taken together, these findings support the idea that CLASH captures proximal mRNAs bound to piRISC via base-pairing interactions.

### piRNA targets exhibit a pattern of discrete peaks in 22G-RNA levels

In *C. elegans*, piRISC recruits RdRP to its targets. Therefore, we wished to examine the pattern of RdRP-dependent 22G-RNA production near CLASH-defined piRNA target sites in both WT and *prg-1* mutant worms. To do this, we plotted 22G-RNA levels within a 40-nt region centered on the piRNA complementary sequences defined by CLASH. The 5’ ends of 22G-RNAs are thought to be formed directly from RdRP initiating at C residues within the target mRNAs. We therefore normalized the 22G-RNA levels initiating at each position to the frequency of C residues within the CLASH-defined targets at each position. Because the CSR-1, and WAGO Argonaute pathway are thought to have opposing functions, resisting and supporting piRNA silencing (Seth et al., 2013; Wedeles et al., 2013), we separately considered predicted piRNA targets within previously defined WAGO and CSR-1 targeted mRNAs (Claycomb et al., 2009; Gu et al., 2009). As a control set, we considered a target region arbitrarily set 100 nts away (within each mRNA) from of the piRNA binding sites identified by CLASH. In WT animals, 22G-RNA levels were much higher for WAGO targets than for CSR-1 targets, as expected (Figures 2A and B, Figures S2A and B). However, piRNA binding sites within both WAGO and CSR-1 targets showed a non-random distribution of 22G-RNA levels across the interval. By contrast, the control regions within the same target mRNAs, but offset from the hybrid sites, exhibited no such patterns (Figure 2G and H). WAGO targets exhibited a strong central peak, and clusters of peaks at either end of the piRNA target sites. To describe these patterns, we refer to the mRNA sequences near the target site as follows: t1 through t30 includes the presumptive binding site (t1 to t21) plus 9 nucleotides 5’ of the target site (t22 to t30). The mRNA region 3’ of the target site consists of nucleotides t–1 through t–11. Strikingly, this analysis revealed a prominent peak in the center of the piRNA complementary region near t12, and smaller peaks centered at t1 and t21 (Figure 2 A). CSR-1 targets exhibited a cluster of much smaller peaks near the 5’ end of the predicted target site, with the largest peak residing in sequences located near t–5 (Figure 2B). The amplitudes of 22G-RNA levels on both the WAGO and CSR-1 targets correlated positively with the predicted free energy of piRNA binding and to a lesser extent with piRNA abundance (Figures 2C-F and S2C-F).

**Figure 2.**
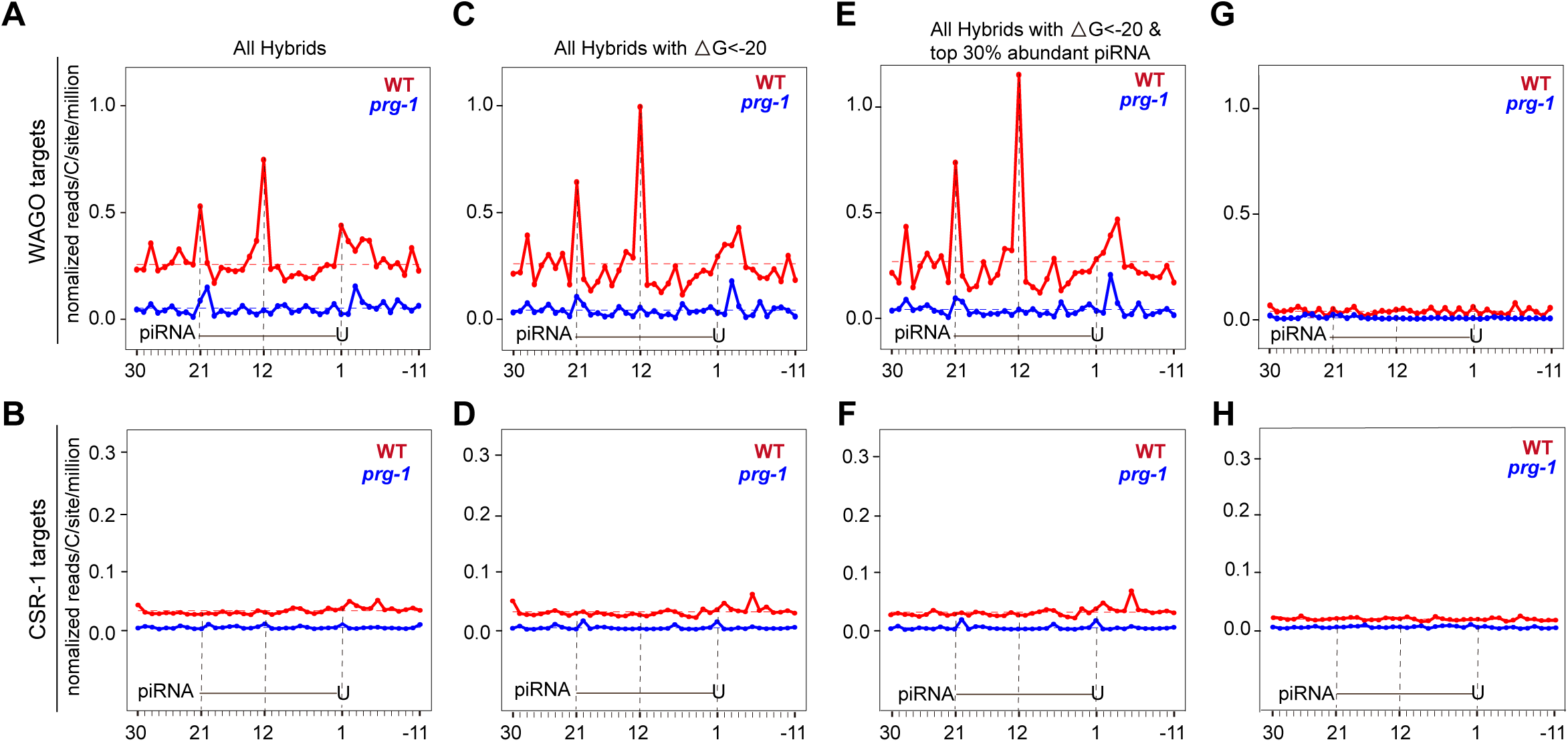
22 G-RNAs peak at the center and ends of piRNA binding sites. **(A-H)** 22G-RNA 5’ ends mapped at single-nucleotide resolution within a 40-nt window around identified piRNA target sites in wild-type (red) or prg-1 mutant (blue) worms. The plots are centered on 21-nt piRNA ± 10 nt shown schematically in each graph. WAGO targets (A, C, E) and CSR-1 targets (B, D, F) were analyzed separately. All hybrids (A and B), hybrids with ΔG < –20 kcal/mol (C and D), hybrids from the top 30% abundant piRNAs with ΔG < –20 kcal/mol (E and F), and control target regions at least 100 nt upstream or downstream of the defined piRNA hybrid sites (G and H). See also Figures S2

The amplitude and position of 22G-RNA peaks differed in *prg-1* mutants. For WAGO targets, the central peak at t12 was completely depleted in *prg-1* mutants, whereas the terminal peaks were reduced. In CSR-1 targets, the prominent peak located at t–5 disappeared, but new peaks at t1, t6, and t21 became evident (Figure 2F). This analysis suggests that PRG-1 influences both the precise position, and the levels of 22G-RNAs on its targets, and that CSR-1 and WAGO targets differ strikingly in their accumulation of 22G-RNAs in response to piRNA targeting (see Discussion).

### Patterns of piRNA targeting

Previous studies have revealed features of Argonaute/small RNA guided targeting, including the importance of "seed" pairing between the target and nucleotides 2–8 of the small RNA guide (Bartel, 2009). To explore patterns of piRNA-mediated targeting we used two independent computational strategies. In the first strategy, we considered the *in silico* predicted folding within a high-confidence group of "ideal" piRNA/target RNA pairs that were identified by at least 5 sequence reads in our combined data sets. To identify preferred base-pairing patterns within this group of hybrids, we applied the Affinity Propagation clustering algorithm (APcluster) (Frey and Dueck, 2007). This analysis revealed a clearly preferred interaction at the seed region and distinct base-pairing patterns at the 3’ supplementary region (Figures 3A and 3B, and Figure S3A). Notably, base-pairing frequencies declined from positions 9 to 13 of the piRNA and increased from positions 14 to 19 (Figure S3B). As expected, these patterns were not enriched in a set of randomized piRNA target RNA pairs (Figure 3A, and Figure S3A).

**Figure 3.**
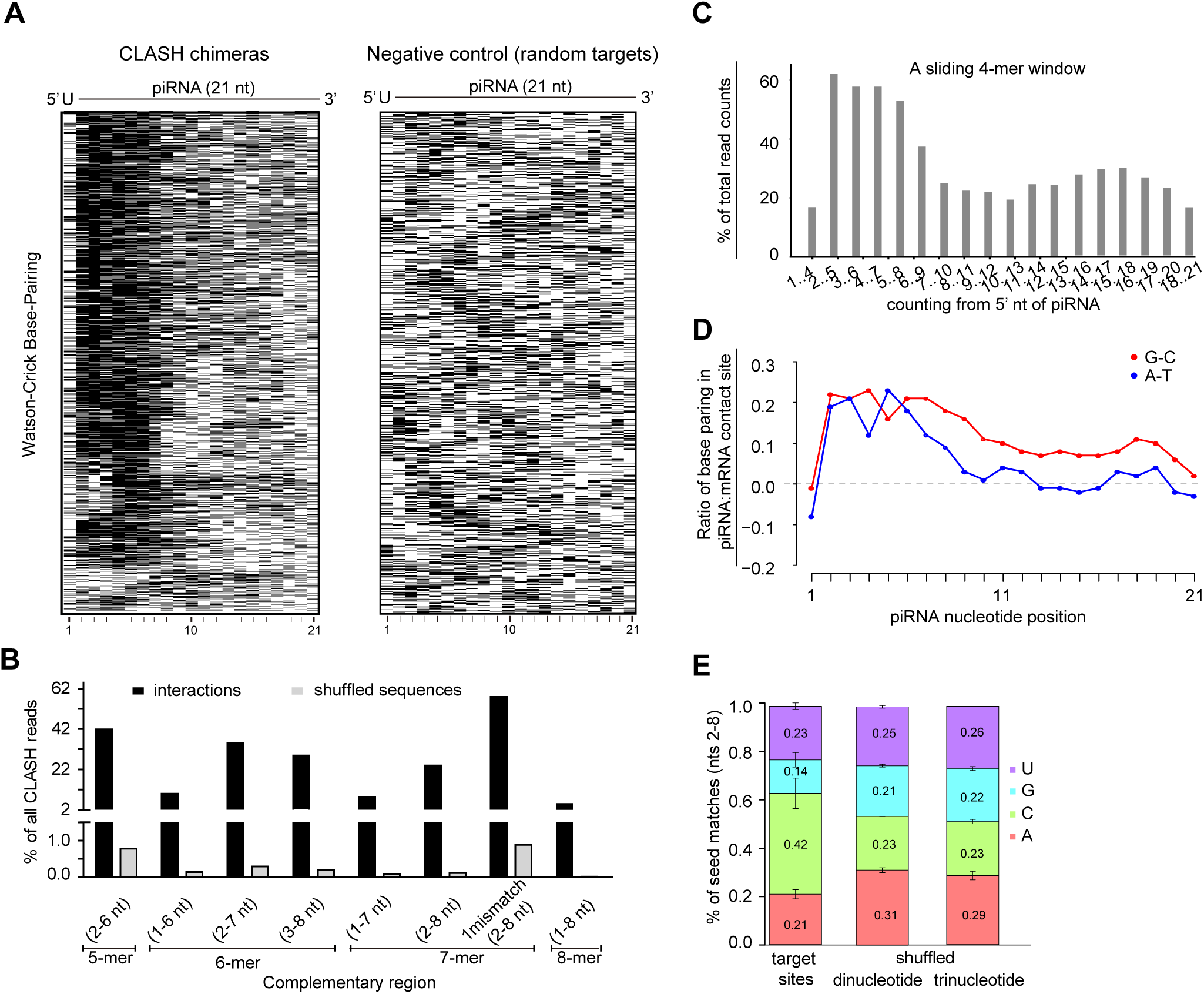
Characterization of piRNA:target base-pairing uncovered by chimeras analysis. **(A)** Heat maps showing Watson-Crick base-paired nucleotides at each piRNA position for all piRNA–mRNA chimeras detected at least 5 times, and for negative control sequences (random target sequences). Black pixels represent pairing. **(B)** Target RNAs were examined for complementarity to positions 1 to 8 of their ligated piRNAs. Approximately 70% of piRNA–target interactions possess the tested complementarities. Complementarities from shuffled dinucleotides of the target sequences served as control. **(C)** Preferential seed and 3’ supplementary pairing between piRNAs and target sites. A 4-mer sliding window search for perfect Watson-Crick base pairing between piRNAs and CLASH chimeras with 50 nt extensions in both directions. **(D)** Ratio of G:C or A:T base pairing in piRNA:mRNA duplexes after deducting ratio calculated from random control. **(E)** T1C is enriched in seed-matched chimeras compared to random target sequences with shuffled dinucleotides and trinucleotides. See also Figures S3

In the second approach, we performed base-pairing analysis using a sliding 4-, 5-, or 7-nt window of piRNA sequence to search for Watson-Crick pairing in each RNA target (Figures 3C, and Figure S3C). Consistent with the analysis in Figure 3A and 3B, this approach revealed a pattern of seed and supplementary pairing, with a distinct drop in pairing from positions 9 to 13 (Figure 3C and Figure S3C). Taken together, these findings suggest that both seed pairing at positions 2 to 8 and supplementary pairing at positions 14 to 19 contribute to piRNA-target RNA binding (Shin et al., 2010; Wee et al., 2012).

To further characterize piRNA–mRNA interactions, we analyzed A:U and G:C base-pair ratios at each position of the piRNA. We found no significant difference between the two base pair ratios within the seed region, but in other regions, we found a bias toward G:C pairing (Figure 3D). Notably, cytosine was strongly over-represented in the target strand immediately 3’ of the seed complement opposite the 5’ u, (defined as target strand position 1 cytosine, or "t1C") (Figure 3E). This preference contrasts with t1A preferred by insect Piwis (Wang et al., 2014). A search for evolutionary conservation using PhyloP scores (Pollard et al., 2010a), failed to reveal a preferential conservation for piRNA-mRNA target sites (Figure S3D).

### Seed and 3’ supplementary pairing are required for target silencing

To determine the importance of base pairing interactions along the length of the piRNA/target mRNA hybrid, we used CRISPR genome editing to systematically mutate positions 2 to 21 of an *anti-gfp* piRNA expressed from the *21ux-1* piRNA locus (Figure 4 A, Figure S4 A-F; Seth et al., submitted). We then assayed the ability of each *21ux-1(anti-gfp)* mutant piRNA to silence a single-copy *cdk-1::gfp* transgene over a time course of up to 8 generations (Figure 4B and C). Strikingly, we found that individual mismatches in the seed region (i.e., m2 to m8) and 3’ supplemental region (i.e., m14 to m21) strongly reduced the ability of *21ux-1(anti-gfp)* to silence *cdk-1::gfp*, but mismatches at the central region (m9 to m13) had a much more mild effect (Figure 4B). By the F2 generation, when fully matched *21ux-1(anti-gfp)* piRNA silences *cdk-1::gfp* by 70% (Figure S4G), mismatches at positions 2 to 8 or 14 to 21 reduced silencing to less than 10% and 25% (respectively) of animals scored (Figure 4B). By contrast, mismatches at positions 9 to 13 reduced silencing activity only slightly, to approximately 50% at the F2 generation. Mismatches at positions 2 or 3 prevented silencing of *cdk-1::gfp* silencing, even after 8 generations, demonstrating that pairing at positions 2 and 3 is essential for piRNA-mediated silencing. Mutants with mismatches at any of the other 18 positions eventually silenced *cdk-1::gfp* over the 8 generation time course (Figure 4C, Figure S4H). Consistent with these findings, we also observed by western blotting after generations 4 and 8 that mutants with mismatches at positions 2 to 8 or 14 to 21 produced much higher levels of CDK-1::GFP protein than did mutants with mismatches at other positions (Figure 4D).

**Figure 4.**
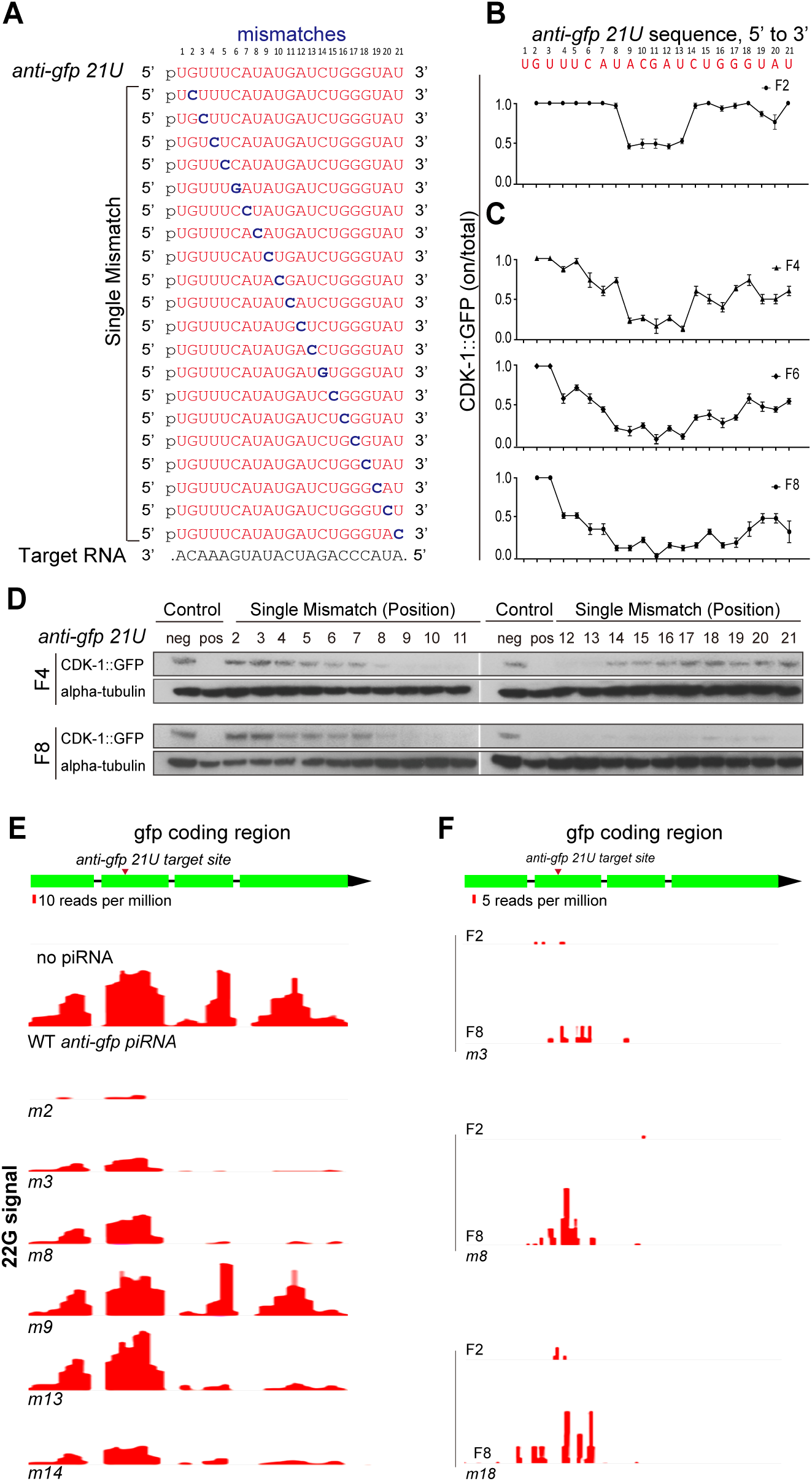
Seed and 3’ supplementary pairing are required for silencing. **(A)** *anti-gfp piRNA* (red) and single-nucleotide mismatches (blue) from positions 2 to 21 on the piRNA target site in *cdk-1::gfp* (black). **(B and C)** Graphs showing the fraction of GFP-positive worms in the presence of *anti-gfp piRNA* with single-nucleotide mismatches at each position. Ten worms (n = 10) were randomly picked for epi-fluorescence analysis at the F2 generation (B) and later generations F4, F6, and F8 (C). Numerals except position 1 denote the order of mutation of *anti-gfp piRNA*. The experiment was performed in triplicate, and data are expressed as means ± s.e.m. **(D)** Western blots showing the levels of CDK-1::GFP protein in F4 and F8 worms (from C) without *anti-gfp piRNA* (negative control), with fully match *anti-gfp piRNA* (positive control), or with single-nucleotide mismatch at the indicated piRNA position. **(E)** Schematic showing density of 22G-RNAs targeting gfp in *cdk-1::gfp* in F4 worms with the indicated single-nucleotide mismatch (m2 = position 2 mismatch, etc.) The positions were randomly chosen and correspond to the 5’, central, and 3’ region of the piRNA. **(F)** Generation-dependent accumulation of 22G-RNA density along the gfp region in *cdk-1::gfp* worms with the denoted mismatch (m3, m8, and m18) at the F2 and F8 generations. Scale bar indicates ten reads per million. See also Figures S4

To further test the importance of pairing in these regions, we selectively mutated positions t3, t15, and t21 of the *anti-gfp* target site in *cdk-1::gfp* mRNA to compensate for *anti-gfp* piRNA mutations in guide-strand positions, g3, g15, and g21, each of which strongly diminished silencing. As expected, in the absence of *21ux-1(anti-gfp)*, these silent mutations did not affect the level of GFP expression (Figure S4I-K). Strikingly, target mRNAs with "re-matching" mutations at t3, t15, and t21 were each rapidly silenced by piRNA strains bearing the corresponding guide mutations (Figure S4K-M). Thus the failure of the g3, g15 and g21 point mutant piRNAs to silence wild-type *cdk-1::gfp* was caused specifically by the mismatches and not by changes in expression or piRISC loading of the mutant piRNAs.

Lastly, we analyzed 22G-RNA induction for several *21ux-1(anti-gfp)* point mutant strains. As expected, we found that 22G-RNA levels correlated with the degree of GFP silencing observed (Figure 4E and F). Overall, these findings confirm the importance of base-pairing within the seed region (nucleotides 2 to 8) and within the 3’ supplemental pairing region (nucleotides 14 to 21) for efficient piRNA targeting.

### Specific piRNA–mRNA interactions suppress endogenous mRNA targets

To investigate how the base-pairing rules defined by our bioinformatics and transgene studies affect targeting of an endogenous mRNA, we edited the *21ux-1* locus, introducing single mismatches into the predicted *21ux-1/xol-1* target duplex (Figure 5A). XOL-1 is a key regulator of dosage compensation and sex determination in early zygotes, and *xol-1* mRNA was recently shown to be regulated by the X chromosome-derived piRNA, *21ux-1* (Tang et al, submitted). Consistent with our findings in the transgene studies, single-nucleotide mismatches within the seed and 3’ supplemental pairing regions, but not within the central region, dramatically increased expression of XOL-1 (Figure 5B). The *21ux-1* mutants with mismatches in the seed and 3’ supplemental pairing regions were phenotypically similar to *21ux-1* null mutants and enhanced the dosage compensation and sex determination phenotypes (decreased brood size and masculinization of hermaphrodites) caused by silencing the X-signal element *sex-1* (Figure 5C and D) (Carmi et al., 1998). Thus, mutating a single nucleotide in *21ux-1* dramatically increases both XOL-1 expression and activity.

**Figure 5.**
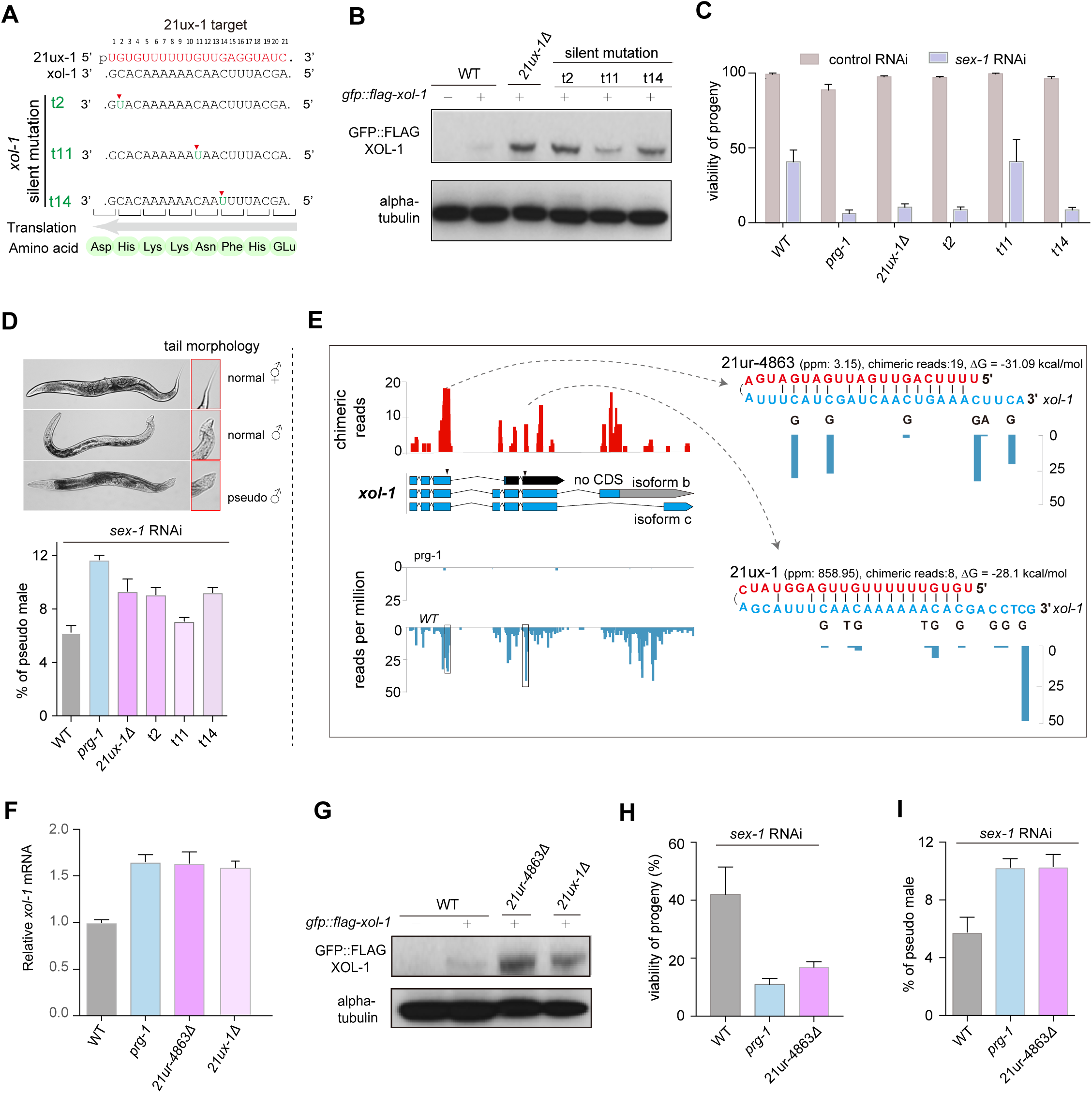
*21ur-4863* and *21ux-1* suppress *xol-1* function in sex determination. **(A)** Diagram showing silent mutations in *xol-1* to create single-nucleotide mismatches with 21ux-1 at positions t2, t11, and t14 (marked by green). **(B)** Western blot analysis of GFP::FLAG::XOL-1 protein expression in N2 worms (negative control, “–”), *gfp::flag::xol-1* transgenic worms (“+”) without *21ux-1* deletion (WT), with *21ux-1* deletion, or with single mismatches (silent mutations) at position t2, t11, or t14. **(C)** Bar graphs showing the percent viable progeny of WT, *prg-1*, *21ux-1* deletion, or *xol-1* single-nucleotide mismatch (t2, t11, t14) worms treated with *sex-1*(RNAi). n > 150 for an experimental group. The experiment was performed in triplicate, and data are expressed as means ± s.e.m. **(D)** Representative DIC images (upper panel) of hermaphrodite, male, and pseudomale worms. Bar graphs (lower panel) showing the percentage of masculinized XX animals in WT, *prg-1*, *21ux-1* deletion, and *xol-1* single-nucleotide mismatch (t2, t11, t14) worms treated with *sex-1*(RNAi). n > 100 for an experimental group. The experiment was performed in triplicate, and data are expressed as means ± s.e.m. **(E)** Distribution of chimeric *xol-1* reads (red) identified by CLASH, and the distribution of *xol-1* 22G-RNAs (blue) in *prg-1* mutant and WT. The sequences and base pairing of *21ur-4863:xol-1* (upper) and *21ux-1:xol-1* (lower) chimeras are shown, and their locations on the *xol-1* gene are indicated by black Inverted triangles. **(F)** Bar graph showing RT-qPCR analysis of *xol-1* mRNA levels in WT, *prg-1*, *21ur-4863* deletion, and *21ux-1* deletion worms. actin mRNA served as the internal control. Data were collected from three independent biological replicates. Error bars represent standard deviation. **(G)** Western blot analysis of GFP::FLAG::XOL-1 (top) levels in WT, *21ur-4863* deletion, and *21ux-1* deletion worms. Alpha-tubulin (bottom) was probed as a loading control. **(H and I)** Bar graphs showing the percent viable (H) and masculinized (I) F1 progeny of WT, *prg-1*, and *21ur-4863* deletion worms treated with *sex-1*(RNAi). n > 500 for an experimental group. The experiment was performed in triplicate, and data are expressed as means ± s.e.m. See also Figures S5

As with most germline mRNAs we found that *xol-1* was targeted by multiple piRNAs. We identified a total of 166 CLASH hybrids containing *xol-1* mRNA sequences fused to 40 different piRNAs (Figure 5E). However, given the importance of *21ux-1* in regulating *xol-1*, and the fact that *21ux-1* is the most abundant piRNA, we were surprised to find that a different piRNA, *21ur-4863*, was recovered in *xol-1* chimeras at a frequency greater than twice that of *21ux-1* chimeras. Specifically, we identified 8 reads with *21ux-1* fused to its *xol-1* target site and 19 reads of *21ur-4863* fused to its *xol-1* target site.

We therefore wished to ask if *21ur-4863* is also important for *xol-1* regulation. Strikingly, deletion of *21ur-4863* resulted in the upregulation of both *xol-1* mRNA and protein levels to a degree similar to that observed in *21ux-1* mutants (Figure 5F and G). Similar to the *21ux-1* mutant, the *21ur-4863* deletion mutant enhanced defects in dosage compensation and sex determination caused by *sex-1(RNAi)*: fewer progeny, masculinization of hermaphrodites, embryonic lethality, and dumpy (Dpy) body morphology (Figure 5H and I, Figure S5 A) (Carmi et al., 1998). Thus, *21u-4863* and *21ux-1* are both required for *xol-1* silencing—neither is sufficient—suggesting that piRNAs function cooperatively to silence *xol-1*.

We also examined the consequences of piRNA targeting on *fbxb-97* and *comt-3*, whose mRNAs are also regulated by PRG-1 (Bagijn et al., 2012; Batista et al., 2008; Gu et al., 2009; Lee et al., 2012). We identified 70 chimeric reads between *21ur-1563* and *fbxb-97* (Figure S5B). *fbxb-97* mRNA levels were upregulated 1.5-fold in a *21ur-1563* deletion mutant and ~8-fold in the *prg-1* mutant (Figure S5C). To analyze piRNA regulation of *comt-3* (Figure S5D), we took the alternative approach of mutating target sequences. We introduced silent mutations into wobble-positions that maintain the *comt-3* open reading frame but disrupt 4 piRNA target sites (Figure S5E). *comt-3* mRNA levels were markedly increased in the *prg-1* mutant and in the *comt-3* quadruple-piRNA target site mutant, but were not elevated in a *comt-3* single-piRNA target site mutant (Figure S5F). COMT-3::FLAG (introduced by CRISPR) was significantly elevated (by 1.5 fold) in the quadruple target site mutant (Figure S5G). Taken together, our findings suggest that individual piRNAs exhibit a range of regulatory effects and that multiple piRNAs cooperatively silence individual targets.

### CLASH analysis reveals competition between the CSR-1 and PRG-1 Argonaute pathways

Previous studies suggested that CSR-1 protects its germline mRNA targets from piRNA-mediated silencing (Lee et al., 2012; Seth et al., 2013; Shirayama et al., 2012; Wedeles et al., 2013). We sought to test whether CSR-1 protects its targets by preventing PRG-1 from binding. To do this, we used an auxin-inducible degradation (AID) system to conditionally deplete CSR-1 in young adult worms (CSR-1^depleted^; Figure S6 A and B) (Zhang et al., 2015a), and then performed CLASH on CSR-1^depleted^ worms in two independent biological replicates. We compared the number of unique piRNA binding sites on CSR-1 targets from CSR-1^depleted^ and wild-type worms. Strikingly, we found that the number of unique piRNA binding sites significantly increased (~2 fold) in the CSR-1^depleted^ worms compared to wild-type (Figure 6A and B, Figure S6C-E). This increase did not result from changes in target mRNA levels, which did not change dramatically during CSR-1 depletion (Figure S6F-H). Increased piRNA targeting is illustrated for *dhc-1*, whose mRNA levels did not appreciably change (~1.5 fold), but whose piRNA targeting was elevated by >3.4-fold in CSR-1^depleted^ worms (Figure 6C). These results suggest that, when CSR-1 is depleted, mRNAs normally targeted by CSR-1 become bound by additional piRNAs.

**Figure 6.**
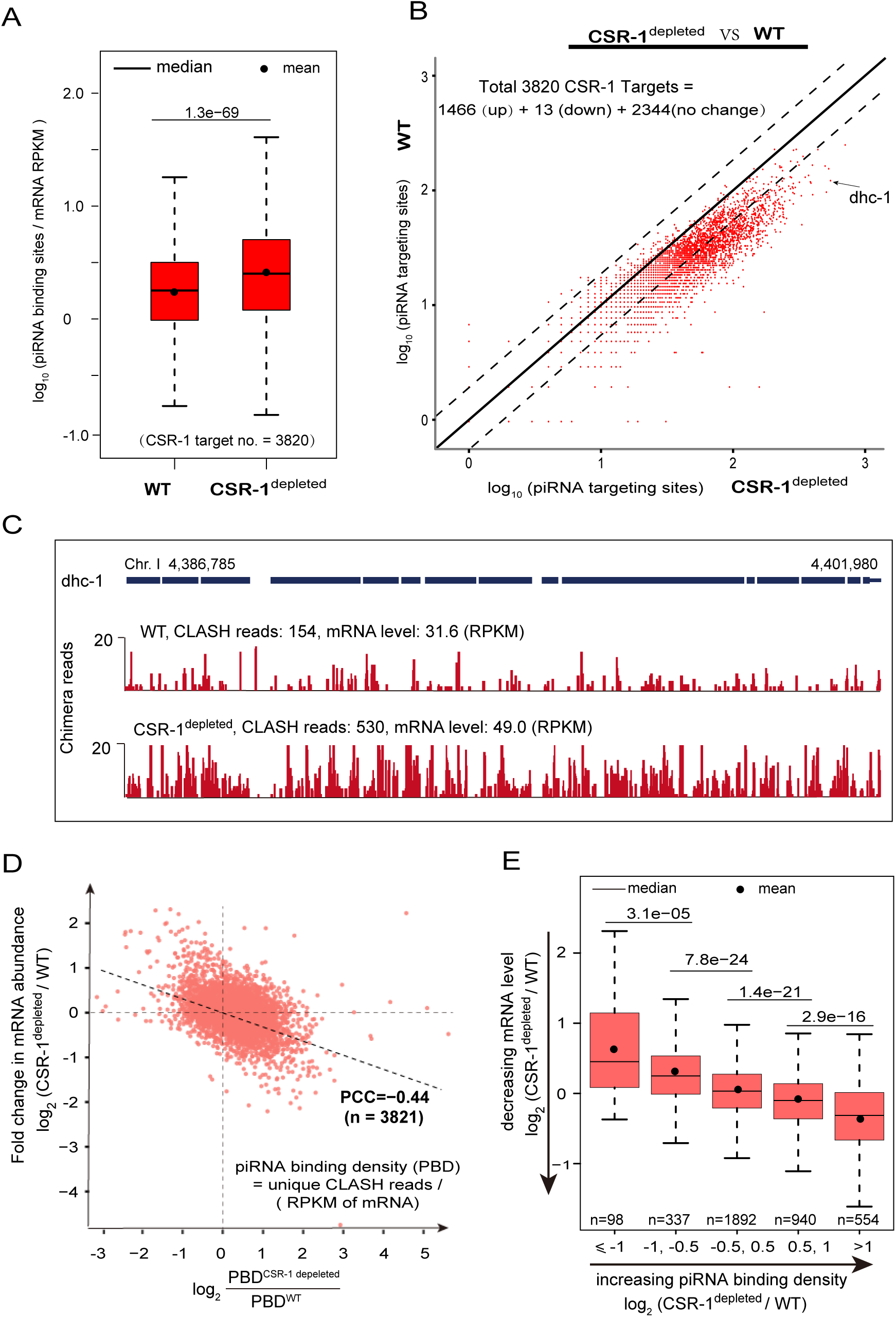
CSR-1 prevents piRNA binding to its targets. **(A)** Box-and-whisker plots of piRNA target sites for CSR-1 targets identified by CLASH in both WT and CSR-1^depleted^ worms. CSR-1 targets amount was normalized using reads per kilobase per million mapped reads (RPKM) in WT and CSR-1^depleted^. Mean and median are indicated separately by black solid line and dot. Outliers removed, and p-values are indicated. **(B)** Scatterplot of unique piRNA binding site counts per gene for each CSR-1 target (n = 3820) in CSR-1^depleted^ versus wild type. For 1466 CSR-1 targets, piRNA binding sites increase >2-fold after depleting CSR-1. A representative target, *dhc-1*, is shown in panel C. **(C)** Genome browser illustrating the distribution of chimeric reads over *dhc-1* in WT and CSR-1^depleted^ worms. **(D)** Scatter plot showing the change in mRNA abundance between WT and CSR-1^depleted^ worms versus the change of piRNA binding density for the 3,821 CSR-1 targets. **(E)** Box plots of the change in mRNA expression levels between WT and CSR-1^depleted^ worms for 5 sets of genes with different levels of piRNA binding site changes. Median and mean are indicated separately by black solid line and dot. Each set is significantly different from the prior one, as indicated by the p-values. See also Figures S6

To determine whether increased piRNA binding correlates with decreased mRNA levels, we plotted the fold change in mRNA abundance (CSR-1^depleted^ / WT) versus the fold change in piRNA-binding density (CSR-1^depleted^ / WT) for all 3,821 CSR-1 targets (Figure 6D) (Claycomb et al., 2009). We observed a negative correlation between increased piRNA-binding density and mRNA abundance in CSR-1^depleted^ worms (r = – 0.44). To clearly visualize this relationship, we split the 3,821 CSR-1 targets into five bins of increasing piRNA binding density and plotted the change in mRNA abundance in CSR-1^depleted^ versus wild type (Figures 6E). This analysis revealed that, as piRNA binding density increases, mRNA abundance decreases. These findings support the idea that CSR-1 functions, at least in part, upstream of PRG-1 to reduce piRNA targeting.

## DISCUSSION

In this study, we took the unbiased approach of directly cross-linking piRNAs to target RNAs *in vivo*. The resulting transcriptome-wide snap-shot of piRNA/target-RNA interactions reveals that all germline mRNAs undergo piRNA surveillance. Our findings are consistent with a model for germline gene regulation wherein mRNAs undergo comprehensive post-transcriptional scanning by Argonaute systems. More than 10,000 distinct piRISCs access hundreds of thousands of target sequences on germline mRNAs. Our finding that binding energy was better correlated with hybrid formation than was piRNA abundance, suggests that, for most piRNAs, piRISC concentration is not limiting. Thus surveillance by piRISC is both transcriptome -wide and remarkably efficient. Perhaps the condensation of piRISC within peri-nuclear nuage creates a local environment that enhances collisions with nascent mRNAs during nuclear export. Considering that substrate release is the slow step in Argonaute surveillance, it will be interesting to determine if factors such as DEAD-box proteins, which also associate with piRISC (Xiol et al., 2014) and are abundantly localized within the nuage environment, function within nuage to facilitate target-release (Figure 7).

**Figure 7.**
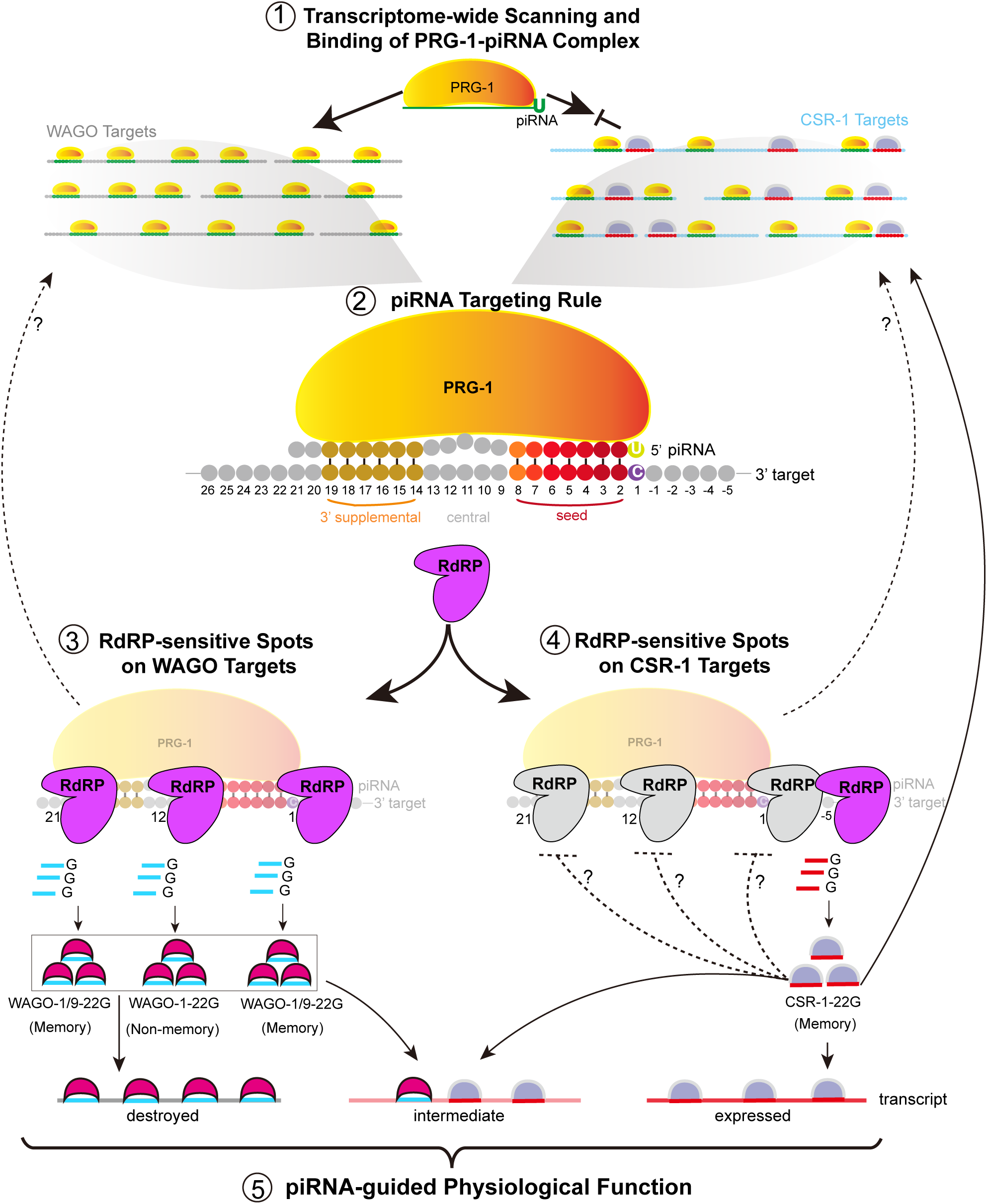
Model for a regulatory landscape of piRNAs in the C. elegans germline

### non-mRNA piRNA interactions

Although mRNA target sites accounted for greater than 90% of CLASH hybrid reads, we also reproducibly identified CLASH reads mapping to a variety of non-coding RNA species (ncRNAs). For example, over 80,000 CLASH reads and hundreds of different piRNAs were mapped to ncRNA hybrids, including sequences from a single region of the 26S rRNA (Figure S1K). Interestingly, this rRNA region is also targeted by WAGO 22G-RNAs that were recently reported to downregulate rRNA levels in response to stress (Zhou et al., 2017).

Our studies also identified many interactions between piRNAs and tRNA species. For example, tRNAGlu(CUC) and 21U-8377 formed highly reproducible chimeras that showed thermodynamically stable base-pairing (Figure S1L). Altogether we identified piRNA tRNA hybrids involving 474 different tRNAs and 1225 different piRNAs. The significance of these findings remains to be determined, but it is intriguing that in *Drosophila* a mutation that leads to accumulation of misprocessed tRNA results in a collapse of Piwi transposon silencing (Molla-Herman et al., 2015; Yamanaka and Siomi, 2015). We also identified hybrids between piRNAs and other ncRNAs including microRNAs and annotated long-noncoding, lncRNAs. The identification of these piRNA interactions provides a new lens through which to explore potential functions and regulation of germline ncRNA species.

### Molecular cross-talk between germline Argonaute pathways

Previous genetic studies have revealed interactions between the Piwi pathway and two Argonaute pathways that propagate epigenetic memories of gene expression states: the WAGO pathway, which targets silenced genes, and the CSR-1 pathway, which targets expressed genes. Targeting by WAGO and CSR-1 Argonautes is readily apparent since both engage 22G-RNAs templated directly from the target RNA by RdRP. Therefore, the comprehensive identification of PRG-1/piRNA target sites affords an opportunity to explore how piRNA targeting correlates with 22G-RNA levels across annotated WAGO and CSR-1 targeted mRNAs.

A striking and unanticipated pattern of 22G-RNA levels emerged from this analysis. On WAGO-targeted mRNAs, piRNA target sites were correlated with three predominant 22G-RNA peaks, one in the center at t12, and one on each side of the targeted site. Interestingly, the central peak at t12 was completely dependent on PRG-1, while the flanking peaks were much less dependent on PRG-1. The flanking peaks that persist in *prg-1* mutants may reflect piRNA-initiated 22G-RNAs that function in WAGO-mediated trans-generational silencing. Consistent with this idea, analyses of data from published WAGO IP experiments indicate that 22G-RNAs at these somewhat *prg-1-*independent flanking sites associate with Argonautes required for propagating piRNA-induced epigenetic silencing (WAGO-1 and WAGO-9) (Figure S2G-J). Interestingly, the strongly *prg-1-*dependent 22G-RNAs generated at t12 associate with WAGO-1 only. Future studies should reveal additional requirements for the PRG-1-dependent 22G-RNAs at t12. For example, it will be interesting to learn why WAGO-1 but not WAGO-9 binds these species and whether their biogenesis depends on PRG-1-dependent mRNA slicing which is predicted to occur between t10 and t11.

Our findings also shed light on the relationship between PRG-1 and CSR-1 targeting. Depletion of CSR-1 resulted in an increase in both unique and total piRNA hybrid reads on mRNAs targeted by CSR-1. These findings are consistent with genetic findings that CSR-1 protects its targets from PRG-1-induced silencing (Seth et al., 2013; Wedeles et al., 2013). Moreover, piRNA target regions on CSR-1 mRNAs exhibit a pattern of 22G-RNA accumulation that is strikingly different from that observed on WAGO-targeted mRNAs. Instead of a central peak and twin flanking peaks as in WAGO targets, a small but reproducible 22G peak, positioned just 5 nt 3’ of the piRNA target site (Figure S2K and L), was evident in CSR-1 mRNAs. Taken together, these findings suggest that CSR-1 protects its targets from piRNA silencing in two ways; first by reducing the frequency of PRG-1 piRISC binding, and second by preventing 22G-RNA accumulation at t12 and flanking regions correlated with WAGO-1 and WAGO-9 targeting.

### Rules governing piRNA Targeting

Our analysis of base-pairing interactions between piRNAs and their targets suggests that animal Piwi- and AGO-clade Argonautes have broadly similar patterns of targeting. As previously described for miRNA RISC, we find that piRISC function strongly depends on pairing in the seed region and to a lesser extent on 3′ supplemental pairing (Shin et al., 2010). The most significant difference we observe is a shift in 3′ supplementary pairing from positions 13 to 16 in miRISC to positions 15 to 18 in PRG-1 piRISC (Grimson et al., 2007), perhaps consistent with structural differences between miRISC and piRISC (Matsumoto et al., 2016a).

In addition to base-pairing interactions, both AGO and Piwi Argonautes make direct contact with their target RNAs, including specific amino acid contacts with the t1 nucleotide. Human AGO2 and insect Piwi proteins (i.e., Siwi and Aubergine) exhibit a strong preference for adenosine at t1 (t1A), which differs from our finding that PRG-1 prefers t1C. This preference for C may help ensure that PRG-1 target sites often have optimal positioning of a C residue that can serve as a start site for RdRP-dependent amplification of 22G-RNAs. A comparison of the region in PRG-1 that corresponds to the t1 binding pocket in other Argonautes suggests a possible structural basis for this discrimination for t1C. Whereas the polar hydrophobic amino acid Thr640 in Siwi and Aubergine is thought to bind t1A (Matsumoto et al., 2016b), the corresponding position in PRG-1 is a non-polar hydrophobic leucine (Figure S3E).

Using a sensitive epigenetic silencing assay, we were able to directly validate the importance of pairing at each position of the seed and 3’ supplemental pairing regions. Silencing was most sensitive to the loss of pairing at positions 2 and 3, suggesting that targeting is initiated by the first half of the seed region. Remarkably, with the exception of positions 9 to 13, which had very little effect on silencing, single nucleotide substitutions at any other location from positions 2 to 8 or 14 to 21 dramatically reduced silencing over the first several generations. Mutants with mismatches in the 3’ supplementary pairing region eventually silenced the target in later generations, but mutants with mismatches in the seed region, especially at g2 and g3, never exhibited full silencing of the target. Thus, seed and 3’ supplementary pairing are of key importance to piRNA targeting. Even single-nucleotide changes dramatically reduced targeting and extended the number of generations required for penetrant silencing.

### The physiology of piRNA targeting

In most animals, Piwi mutants are completely sterile, likely due at least in part, to loss of transposon regulation. In worms, most transposons appear to be silenced by epigenetic mechanisms—i.e., WAGO 22G-RNAs and heterochromatin pathways—that maintain transgenerational silencing downstream of PRG-1 (Ashe et al., 2012; Bagijn et al., 2012; Gu et al., 2009; Lee et al., 2012; Shirayama et al., 2012). This additional layer of epigenetic silencing may explain why *prg-1* mutants exhibit relatively minor transposon activation and fertility defects during early generations, but exhibit declining fertility over multiple generations (i.e., a mortal germline phenotype) (Simon et al., 2014).

PRG-1 is nevertheless constantly required to maintain silencing at some loci. Transgenes exposed to both positive (i.e., CSR-1-dependent) and negative (i.e., piRNA-dependent) signals can achieve a balanced state of regulation, where PRG-1 targeting becomes essential to maintain silencing (Seth et al., submitted). At least a few hundred endogenous mRNAs are significantly up-regulated in *prg-1* mutants, with a concomitant loss of robust 22G-RNAs levels. One such gene, *xol-1*, is silenced in the hermaphrodite germline by an X-chromosome expressed piRNA, *21ux-1* (Tang et al., and Meetu et al., submitted). Silencing of *xol-1* ensures that hermaphrodite offspring respond robustly to signals that initiate dosage compensation and sex determination in the early embryo. Although *21ux-1* is by far the most abundant piRNA species, a piRNA with average abundance (*21ur-4863*) binds *xol-1* more efficiently based on the frequency of CLASH hybrid identification. 21ur-4863 is predicted to bind *xol-1* with higher binding energy than predicted for *21ux-1*, highlighting the importance of binding energy rather than abundance in driving piRNA targeting. Surprisingly, both *21ur-4863* and *21ux-1* are required to maintain *xol-1* silencing, suggesting that they—and perhaps other—piRNAs cooperatively silence *xol-1*. Remarkably, even though multiple piRNAs regulate *xol-1*, changing a single nucleotide within the seed or 3’ supplementary pairing regions of *21ux-1* can disrupt silencing of *xol-1* and thus affect the regulation of dosage compensation and sex determination.

In summary, our findings show that piRNAs target the entire germline transcriptome. Together with findings from previous and parallel studies our findings also suggest that piRNAs are remarkably versatile in their control of gene expression. piRNAs can act decisively in one generation to initiate epigenetic silencing that persists for multiple generations without need for further piRNA targeting. piRNAs can act cooperatively to silence germline mRNAs (e.g., *xol-1*) that would otherwise reactivate in each generation. And finally, piRNAs can act gradually, over multiple generations, to progressively silence a germline mRNA. Understanding how piRNAs achieve these nuanced modes and tempos of regulation may shed light on whole new vistas of post-transcriptional and epigenetic regulation in animal germlines.

## EXPERIMENTAL PROCEDURES

### *C. elegans* Strains

Strains in this study were derived from the Bristol N2 background, and cultured on NGM agar plates with OP50 *E. coli* at 20°C (Brenner, 1974). All the transgene insertions are generated by using a Mos transposon-based strategy (Frøkjaer-Jensen et al., 2008; Frøkjær-Jensen et al., 2012; Shirayama et al., 2012).

### CRISPR/Cas9 genome editing

The *gfp::tev::flag::prg-1*, *21ux-1* deletion, and *aid::csr-1* strains were generated using an unc-22 co-CRISPR strategy, in which F1 animals with a twitching phenotype (unc-22 mutant) are enriched for CRISPR/Cas9-mediated genome editing events (Kim et al., 2014). Guide RNA vectors were constructed as described (Arribere et al., 2014). For generating transgene strains, another strategy is injecting preassembled CRISPR/Cas9 ribonucleoprotein complexes using *rol-6* as a co-injection marker (Paix et al., 2015). Genome-edited animals were identified among F1 rollers. The online tool CRISPR DESIGN (http://crispr.mit.edu) was used to design guide RNAs. All strains are listed in Supplemental Table 1.

### Small RNA Library Preparation and analysis

Total RNA was extracted from synchronous adult worms using TRI Reagent (Molecular Research Center), according to the manufacturer’s instructions. Small RNAs were extracted from total RNA using mirVana miRNA Isolation Kit (Thermo Fisher Scientific). Samples were pretreated with homemade, recombinant 5 ′ polyphosphatase PIR-1 to remove the Ɣ and β phosphates from triphosphorylated 5′ ends of 22G-RNAs (Gu et al., 2009). The resulting 5′ -monophosphorylated small RNAs were ligated to a 3 ′ adapter (5 ′ rAppAGATCGGAAGAGCACACGTCTGAACTCCAGTCA/3ddC/3 ′; IDT) using T4 RNA ligase 2. The ligation products were then ligated to 5 ′ adapter (rArCrArCrUrCrUrUrUrCrCrCrUrArCrArCrGrArCrGrCrUrCrUrUrCrCrGrArUrCrU) was using T4 RNA ligase 1. Ligation products with 5′ and 3′ linkers were reverse transcribed using SuperScript II. The cDNAs were amplified by PCR, and the libraries were sequenced on an Illumina HiSeq platform at the UMass Medical School Deep Sequencing Core Facility.

Small RNA sequencing data were analyzed as described (Lee et al., 2012). Briefly, both 5’ and 3’ adapter sequences were trimmed using a custom Perl script. Sequencing reads were sorted into different bins according to barcode sequences, and 21-23nts reads were mapped to the C. elegans genome (Wormbase release WS230) allowing no more than 2 mismatches. To account for variation in sequencing depth between samples, each read was normalized to the total number of reads. Normalized counts were visually observed in the UCSC genome browser. All scripts are available upon request.

### mRNA Library Preparation

Total RNA from worms was extracted using TRI Reagent (MRC) and ethanol precipitation. RNA samples were processed as described (Zhang et al., 2012). Briefly, ribosomal RNAs were depleted from 4 µg total RNA using the RiboMinus Eukaryote Kit (Life Technologies, A1083708), and the rRNA-depleted samples were treated with Turbo DNase (Ambion) for 30 min at 37°C. RNAs >200 nt were enriched using RNA Clean & Concentrator-5 (Zymo Research, R1015), fragmented, and reverse transcribed into cDNA using oligo-dT and SuperScript II. The first-strand cDNA was ligated to 5′ and 3′ adapters using T4 DNA ligase (600 U/µL, Enzymatics Inc, L603-HC-L) for 30 min at 25°C. The library was amplified with a barcoded PCR primer. Deep sequencing was performed on an Illumina HiSeq 2000.

### Immunoprecipitation and RNA Isolation

200,000 synchronous adult worms were homogenized in a FastPrep-24 benchtop homogenizer (MP Biomedicals). Immunoprecipitation was performed as described (Tang et al., 2016). In brief, worm extracts were cleared by centrifugation for 10 min at 14,000 × g. Lysates (20 mg total protein) were incubated with anti-FLAG M2 magnetic beads for 1.5 h at 4 °C on a rotator. Beads were washed 3 times with IP buffer containing protease inhibitors for 10 min each wash, and then washed once with wash buffer containing 50 mM Tris·Cl (pH 7.5), 2 mM magnesium acetate, and 150 mM NaCl. Immune complexes were treated with 10 mg/ml proteinase K in 2.5% (w/v) SDS, 200 mM EDTA, 100 mM Tris-HCl (pH 7.5) for 10 min at 50°C. RNAs were extracted with acidic phenol/chloroform (low PH) and precipitated with ethanol. Isolated RNA was treated with PIR-1 and subjected to small RNA cloning.

### RT-qPCR

Total RNA was treated with Turbo DNase (Ambion), extracted with TRI Reagent (MRC), and precipitated with ethanol. 500 ng of RNA was used as a template for first-strand cDNA synthesis using random hexamers and SuperScript II Reverse Transcriptase (Invitrogen) according to the manufacturer’s instructions. cDNA synthesis was performed in triplicate. Real-time PCR was conducted using Power SYBR Green PCR Master Mix (Applied Biosystems) on the Bio-Rad CFX96 Real Time PCR Detection System. Primer sets I (CAGCTGGAAATTACCGAGGA and GTTGGCCATGGAACAGGTAG), II (AGGTGATGCAACATACGGAA and CGAGAAGCATTGAACACCAT), III (CATGGCCAACACTTGTCACT and GCACGTGTCTTGTAGTTCCC), IV (TCAAAGATGACGGGAACTAC and GCTTCCATCTTCAATGTTGT) were used to amplify *gfp* transcript. Primers AGCTTCTTCGAGATGCGTTC and CTTGTCGCACACGGTTCTTG were used to amplify *xol-1* transcript. Primers TCGGAGTTCCGATCATCTCG and CAGGGTGACAGCTCTATCGT were used to amplify *comt-3* transcript. Primers GGCCCAATCCAAGAGAGGTATCC and GGGCAACACGAAGCTCATTGTA were used to detect *act-3* mRNA. Error bars in the graph indicate the standard deviation (SD) in all statistical analysis.

### Western Blot Analysis

Protein lysates were prepared from synchronous populations of young adult or gravid adult worms. Proteins (50 µg) were separated on precast denaturing polyacrylamide gels (Thermo Fisher Scientific) and transferred to PVDF membranes (Bio-Rad) using a Trans-Blot Turbo Transfer System (Bio-Rad). Membranes were blocked and probed with primary antibodies: monoclonal anti-FLAG (Sigma, M8823), monoclonal anti-GFP (Roche, 11814460001), monoclonal anti-TUBULIN ALPHA (AbD Serotec, MCA77G), and anti-mini-AID-tag (MBL International, M214-3). Primary antibodies were detected using HRP-linked secondary antibodies: donkey anti-rat IgG(Jackson ImmunoResearch, 712–035-150) or goat anti-mouse (Thermo Fisher Scientific, 62–6520).

### RNAi

RNAi was performed by feeding worms *E. coli* strain HT115(DE3) transformed with the control vector L4440 or a gene-targeting construct from the *C. elegans* RNAi Collection (Kamath and Ahringer, 2003). Bacteria were grown overnight in LB with antibiotics at 37°C. NGM plates containing 1 mM isopropyl β-d-thiogalactoside and 100 µg/ml ampicillin were seeded with the overnight culture (100 µl per plate) and incubated for 24 hours at 25°C. L1 larvae were placed on RNAi plates at room temperature and the phenotypes of the adults and their F1 progeny were scored.

### Microscopy

Transgenic worms expressing mCherry or GFP were anaesthetized in 0.1 mM levamisole (Sigma, 16595–80-5) on glass slides with 10-mm superfrost circles (Thermo Scientific, 3032) and imaged immediately. Epi-fluorescence and differential interference contrast microscopy were performed in a Zeiss Axioplan2 microscope. Images were captured and processed using Zeiss Axiovision software.

### Auxin-inducible Depletion of CSR-1

Auxin treatment was performed as described (Zhang et al., 2015a). 100 mM indole-3-acetic acid (IAA; Alfa Aesar, 10171307) was prepared in ethanol and stored for up to one month at 4°C. NGM plates containing 500 µM IAA (prepared by adding IAA to NGM agar at ~50°C) were seeded with fresh concentrated OP50 and incubated at RT for 48 hours. IAA plates stored at 4 °C were warmed to room temperature for 1 hour before use. *aid::csr-1* (WM509) or wild-type worms were placed on IAA plates as L1 larvae and grown to the young adult stage at 20°C.

### Viability and Pseudomale Development

Ten gravid hermaphrodites (WT or mutant) were picked onto individual plates, and transferred to new plates every 12 hours. The number of embryos and hatched larvae were counted at each transfer, and the plates were summed to determine the total number of progeny per hermaphrodite. Progeny were allowed to develop at room temperature for several days, and the number and phenotypes of F1 adult animals were scored at room temperature. Viability is expressed as the percentage of F1 progeny that develop into adults. Pseudomales were identified as F1 adult hermaphrodites that develop male-like tails. *sex-1*(RNAi) was used to enhance the sex-determination defect (masculinization) caused by *xol-1* gain of function.

### PRG-1 CLASH Protocol

#### Worm growth, UV-crosslinking, and worm lysis

L1 worms were grown on large NGM plates with concentrated OP50. Synchronous adult worms (~1,000,000) were collected and transferred to 50 large NGM plates without OP50. Plates were placed on ice in a Stratalinker 1800 (Stratagene) and worms were irradiated with 254-nm light at 1.0 J/cm^2^ to crosslink protein and RNA.

Irradiated worms were homogenized in cooled lysis buffer composed of 20 mM HEPES-KOH (pH 7.5), 125 mM citrate sodium, 2 mM MgCl2, 1 mM DTT, 0.5% (v/v) Triton X-100, 0.1% (v/v) Tween 20, 0.04% protease inhibitors (Complete Mini EDTA-free; Roche) using the FastPrep-24 homogenizer. Lysates were centrifuged at 14,000 rcf. (x g) for 10 min at 4°C, and supernatants were treated with RNase T1 (3 U/µL) for 15 min at room temperature on a rotator.

#### GFP pull-down and TEV cleavage

Worm lysates were incubated with GBP beads for 2 hrs at 4°C, and then beads were washed three times with washing buffer I (50 mM HEPES-KOH, pH 7.5, 300 mM KCl, 0.05% NP-40, 0.5 mM DTT, 0.04% protease inhibitors) for 5 min each, twice in washing buffer II (50 mM HEPES-KOH, pH 7.5, 500 mM KCl, 0.05% NP-40, 0.5 mM DTT, 0.04% protease inhibitors) for 5 min each, and once in TEV buffer (50 mM Tris-HCl pH 8.0, 0.5 mM EDTA, 1 mM DTT, 2 mM MgOAC) for 5 min. Beads were suspended in 100 µL TEV buffer, and 8 µL AcTEV Protease was added to cleave TEV site between the GFP and FLAG tags. The cleavage reaction was incubated for 2 hrs at room temperature, releasing FLAG::PRG-1/RNA complexes into solution.

#### Purification of FLAG::PRG-1/RNA complexes

To capture FLAG::PRG-1/RNA complexes, anti-FLAG antibodies were incubated with Protein G dynabeads (Life Technologies) for 30 min at room temperature. Beads were washed three times with washing buffer I for 5 min each, twice with lysis buffer containing 0.5% BSA for 10 min each, and twice with lysis buffer containing 0.01% BSA for 10 min each. FLAG::PRG-1/RNA complexes were captured by incubating the TEV-released complexes with washed Dynabeads in TEV buffer containing 2% RNase inhibitor for 1 h at 4°C.

#### 5’ end phosphorylation and intramolecular ligation

FLAG::PRG-1/RNA immune complexes were washed once with 1× PBS, 2% Empigen (PMPG buffer: Sigma, 30326), once with 5× PBS, 2% Empigen, and once with 1× PNK buffer. FLAG::PRG-1/RNA complexes were treated with 0.5 U/µl T4 Polynucleotide Kinase in the PNK buffer (1 mM DTT, 1 mM rATP, 1 U/µL RNase inhibitor, 0.005 U/uL Turbo DNase) for 30 min at room temperature, and then washed once with 1× PMPG buffer, once with 5× PMPG buffer, and once with 1× PNK buffer. FLAG::PRG-1-bound, RNAs molecules were ligated using 2 U/µl T4 RNA ligase in 1x ligase buffer (containing 1 mM rATP, 1 U/uL RNase inhibitor) without DTT for 2 hrs at 16°C on a rotator. The ligation mixture was washed once with 1× PMPG buffer, once with 5× PMPG buffer, and once with 1× PNK buffer.

#### RNA dephosphorylation and ^32^P-labeled 3’ linker ligation

The FLAG::PRG-1/RNA complexes on Dynabeads were incubated with 0.1 U/µL FastAP Thermosensitive Alkaline Phosphatase (Life Technologies, EF0652) in 1× FastAP buffer containing 1 U/µl RNase inhibitor for 30 min at room temperature on a rotator. Complexes were then washed once each with 1× PMPG buffer, 5× PMPG buffer, 1× PNK buffer, and 1× RNA ligase buffer without DTT. Dephosphorylated RNA was incubated with 33 uM 3’ linker (～10% ^32^P-labeled) in 1× ligation buffer containing 2.5 U/µL RNA ligase, 1 mM rATP, 2.5% DMSO, 15% PEG 8000, and 1 U/µL Rnase inhibitor for 2 hrs at 16°C. Beads were washed once each with 1× PMPG buffer, 5× PMPG buffer, and 1× PNK buffer.

#### Elution of FLAG::PRG-1/RNA complexes, SDS-PAGE, and transfer to nitrocellulose

FLAG::PRG-1/RNA complexes were eluted in 1× SDS-PAGE loading buffer for 10 min at 70°C. Duplicate samples (one for western blot analysis one to recover complexes with ligated linkers) were separated on a NuPAGE 4–12% Bis-Tris protein gel (Life Technologies, NP0321) using NuPAGE SDS-MOPS running buffer (Life Technologies, NP0001) in a cold room. FLAG::PRG-1/RNA complexes were transferred to pre-cut nitrocellulose blotting membrane (Invitrogen, LC2000) in a wet-transfer tank with NuPAGE transfer buffer containing 10% methanol, overnight at 30 V. The membrane was cut into two parts along a marker. One part was analyzed by western blot analysis using anti-FLAG antibody as described above. The other part of the membrane was air dried and exposed to film (Sigma, Z350370) for 2 hrs at –80°C. The developed film was aligned to the membrane and ^32^P-labeled region of the membrane containing FLAG::PRG-1/RNA complexes were excised.

#### Proteinase K treatment and RNA extraction

Membrane slices were treated with 50 µl of proteinase K (New England Biolabs, P8107S) in proteinase K buffer (100 mM Tris-HCl, pH 7.4, 50 mM NaCl, 10 mM EDTA) for 20 min at 37°C, and then 42% Urea/Proteinase K buffer was added into sample for an additional 20 min at 37°C. RNA was extracted with phenol/chloroform/isoamyl alcohol, and then precipitated with ethanol/isopropanol and 5 µg GlycoBlue (Ambion, AM9516), overnight at –80°C. RNA pellets were washed with 70% cold ethanol and dissolved in RNase-free water.

#### 5’ phosphorylation, RNA isolation, and 5’ linker ligation

RNAs were incubated with 0.5 U/µL T4 Polynucleotide Kinase in 1× PNK buffer (1 mM ATP, 1 U/µL RNase inhibitor) for 30 min at 37°C. RNA was purified using an RNA Clean & Concentrator Kit (Zymo Research, R1016) according to the manufacturer’s instructions. RNA was ligated to 10 uM 5’ linker using 2 U/µL RNA ligase in 1× ligation buffer containing 1 mM ATP, 2.5% DMSO, 8% PEG 8000, 1 U/µL RNase inhibitor for 3 hrs at 16°C.

#### RNA extraction, reverse transcription, and library construction

5’-ligation products were purified using RNA Clean & Concentrator Kit, and converted to first-strand cDNA using primer (GTGACTGGAGTTCAGACGTGTGCTCTTCCGATCT) with Superscript III Reverse Transcriptase (Life Technologies, 18080–093), according to the manufacturer’s instructions. cDNA was amplified using linker primers and Q5 High-Fidelity DNA Polymerase (New England Biolabs, M0491S), according to the manufacturer’s instructions. PCR products were separated on a 12% polyacrylamide gel in 1× TBE buffer and stained with ethidium bromide. Gel slices containing amplified libraries (160 to 190 bp) were excised and the library was extracted using a Gel Extraction Kit with columns (QIAGEN).

### Bioinformatic Analysis of P-CLASH Data

#### Pipeline for mapping CLASH hybrids

CLASH chimeras containing full-length piRNA sequences were initially selected from the library using Bowtie, version 0.12.9 (Langmead et al., 2009). piRNAs were then trimmed from the CLASH chimeras, and the remaining sequences were mapped to the *C. elegans* genome (Wormbase WS230) (Yook et al., 2012). These candidate piRNA-target sequences were then classified based on annotations using a customized Perl script and BEDtools (version 2.25.0) (Quinlan and Hall, 2010). As annotation is missing in this release, lincRNAs and Tc1 and Tc3 transposons were manually annotated by mapping their sequence to the genome using BLAT (version 35x1) (Kent, 2002).

#### piRNA-target duplex prediction, CLASH chimera refinement, and clustering

piRNA-target duplexes and changes in Gibbs free energy (∆G) were calculated using RNAfold in Vienna RNA Package (version 2.3.5) (Lorenz et al., 2011). Based on the initial alignment, additional bases were added to the 5’ or 3’ ends of trimmed target sequences, based on longer CLASH chimeras or on the reference genome. The duplex prediction and ΔG were recalculated for these refined target sites.

Matching relationships for piRNA-mRNA chimeras detected at least 5 times were summarized as follows Alignments from predicted duplexes were converted into matches, mismatches, and internal bulges using BEAR encoder (Mattei et al., 2014). The piRNA side of the duplex was clustered using affinity propagation (Frey and Dueck, 2007) with a preference score of 5000. The similarity score for clustering was defined as a negative edit distance between duplexes, ranging from 0 to –21. The cluster plot was then generated using deepTools (version 2.5.1) (Ramírez et al., 2016).

#### Conservation

Pre-calculated phyloP scores for the C. elegans genome (version WS220) based on 7 Caenorhabditis species were downloaded from UCSC browser (Kent et al., 2002; Pollard et al., 2010b). phyloP scores are only available for 9400 coding genes. The piRNA target sites on these genes were separated into 3 groups according to their starting codon position. Then, the average phyloP scores for genomic positions that are on, around, or off the target site were calculated.

#### RNA-seq data processing and differentially expressed gene analysis

Adapter sequences and low-quality reads were removed using Trimmomatic, version 0.33 (Bolger et al., 2014). The short reads that passed quality control were mapped to the genome using STAR, version 2.4.2a (Dobin et al., 2013). The read counts for each gene with uniquely mapped strand-specific reads were calculated using Samtools, version 0.1.19 (Li et al., 2009) and HTseq, version 0.7.2 (Anders et al., 2015).

#### 22 G-RNAs around the piRNA target sites

Adapter sequences and low-quality reads were removed using Trimmomatic (version 0.33). The short reads were then mapped to the genome using STAR (version 2.4.2a) and normalized to sequencing depth. Short reads 21- to 23-bp long with 5’ G were defined as 22 G-RNAs. Their 5’-end signals were aggregated for each position around the target sites and normalized to the number of transcriptomic C at these positions.

#### Normalization of CLASH counts

The number of unique CLASH hybrids in each library was normalized to the mean tRNA hybrid counts, using their trimmed mean as the normalization factor.

#### Scatter plots

Scatter plots were generated using ggplot2 (Wickham, 2009).

#### CSR-1 and WAGO targets

The 4192 CSR-1 target genes and 1118 WAGO target genes were previously defined (Claycomb et al., 2009; Gu et al., 2009).

#### Soma-specific genes

modENCODE RNA-Seq data has defined 2423 soma-specific genes expressed in young adult worms (Li et al., 2014b).

#### piRNA target sequence shuffling

piRNA target sequences were shuffled while maintaining their di-nucleotide frequencies using uShuffle (Jiang et al., 2008).

#### Data availability

The authors declare that the raw data are available within the paper and its supplementary information files using the accession codes BioProject: SUB3396

## Author Contributions

Conceptualization, E. Z. and C. C. M.; Investigation, E. Z., H. C., A. O., S. T., M. S., W. T., Y. D., S. D., D. C., Z. W. and C. C. M.; Writing-Original draft, E. Z. and C. C. M.; Writing-Review & Editing, E. Z. and C. C. M.; Supervision, C. C. M and Z. W.

## Acknowledgement

We thank P. Zamore and V. Ambros for suggestions; members of Mello and Ambros labs for discussions, D. Conte for comments and edits on the text; W. Gu for providing the reagents for preparing library; M. Carmell for proofreading; E. Kittler, D. Wilmot and the Umass Deep Sequencing Core for offering high-throughput sequencing; Caenorhabditis Genetics Center (CGC) for providing strain. This work was supported by an NIH grant (HD078253) to Z. W.; and NIH grants (GM058800 and HD078253) to C. C. M. C. C. M. is a Howard Hughes Medical Institute Investigator.

---

**Figure S1. Two independent P-CLASH Data Sets, Related to Figure 1**

**(A)** Fluorescence micrographs showing the endogenous expression of GFP::TEV::FLAG::PRG-1 in C. elegans germline. Schematics of *gfp::tev::flag::prg-1* (*gtf::prg-1*) was shown on the top. prg-1 was tagged with *gfp::tev::flag* by CRISPER-55 CAS9 technique. mCherry::PGL-1 (red) served as the P granule marker.

**(B)** Image of *gfp::cdk-1* reporter in the wild-type (WT), *prg-1(tm872)* and *gtf::prg-1* strains. *gfp::cdk-1* construct on the top.

**(C and D)** The graphs present the number of reads mapped to each identified piRNA (C) and mRNA (D) in one experiment relative to the other one. r – Pearson’s correlation coefficient.

**(E and F)** Venn diagram summarizing the overlap in the number of piRNAs (E) and mRNA (F) in two replicates.

**(G and H)** Distribution of all piRNA interactions among various types of RNAs in two replicates. mRNA are the main piRNA targets and represent more than 70%.

**(I)** Correlation of piRNA abundance between total piRNA abundance (input) (y-axis) and P-CLASH recovered piRNA (x-axis).

**(J)** Histogram distribution of unique piRNA CLASH hybrids. The negative value means no nucleotides are added to the target sequence. All CLASH hybrids with values >= 11 or <= – 11 are summed up. Dark blue indicates the ligation events happened at the 3’ end of piRNA and red indicates those CLASH hybrids ligate at the 5’ end of piRNA.

**(K)** Distribution of chimeric *rrn-2.1* and *rrn-3.1* reads (green) identified by CLASH and genomic locus on the top (blue).

**(L)** Putative interactions between piRNA and tRNA. *21ur-8377* reproducibly bind to the same region of tRNA-Glu(CUC), marked blue on the tRNA structure (chrIII. ZK783.t1).

**Figure S2. 22G-RNAs signal in the piRNA binding region, Related to Figure 2**

**(A-F)** 22G-RNAs peak at the center and ends of piRNA binding sites in another dataset, similar to Figure 2.

**(G and H)** Distribution of 22G-RNA signal in input versus WAGO-1 IP for WAGO targets (G), and for CSR-1 targets (H).

**(I and J)** Distribution of 22G-RNA signal in input versus WAGO-9 IP for WAGO targets (I), and for CSR-1 targets (J).

**(K and L)** Distribution of 22G-RNA signal in input versus CSR-1 IP for CSR-1 targets (K), and for WAGO targets (L).

**Figure S3. Related to Figure 3.**

**(A)** Positions of base-paired nucleotides in piRNAs for all piRNA:mRNA interactions. piRNA:targets duplex structure predictions were calculated using RNAfold in Vienna RNA Package, and clustering was analyzed using APcluster. Shuffled interactions (targets are swapped between piRNAs) served as random control. Base-paired nucleotides are shown in piRNAs length.

**(B)** Hybridization profile showing all interactions. The predicted frequency of a piRNA position to be base paired is plotted along the piRNA.

**(C)** Preferential seed and 3’ supplementary pairing between piRNAs and target sites. Similar analysis to Figure 2C, a 5-mer and 7-mer sliding window search are performed.

**(D)** Average mRNA conservation level on the piRNA-miRNA contact sites. C. elegans phyloP score starting from each codon position at the target sites. See also method.

**(E)** Alignment of Multiple Sequence for the Argonaute Proteins. PRG-1 sequence is compared with published t1A binding pocket data (Matsumoto et al., 2016b). The figure was prepared using ESPript3 (http://espript.ibcp.fr/ESPript/ESPript).

**Figure S4. *anti-gfp piRNA* silences its target *cdk-1::gfp* in C. elegans, Related to Figure 4.**

**(A)** Generation of CRISPR-Cas9-mediated *anti-gfp piRNA* knock-in strain.

**(B)** CDK-1::GFP expression in *cdk-1::gfp*, *cdk-1::gfp; anti-gfp piRNA* and *cdk-1::gfp; anti-gfp piRNA; rde-3* worms. Images of CDK-1::GFP fluorescence signals in the resulting strains are shown. Germline nuclei that express CDK-1::GFP are denoted by arrowheads. The dashed lines outline the position of germline. Bright signals outside of the germline are from gut granule autofluorescence.

**(C)** RT-qPCR analysis of *cdk-1::gfp* mRNA level in N2, *cdk-1::gfp*, *cdk-1::gfp; anti-gfp piRNA* and *cdk-1::gfp; anti-gfp piRNA; rde-3* worms. Schematics of the single-copy transgene *cdk-1::gfp*. Red arrowhead indicates target site of *anti-gfp piRNA.* The black line beneath the schematics represent the location of PCR product.

**(D)** Western blot analysis of the Change of CDK-1::GFP in N2, cdk-1::gfp, cdk-1::gfp; *anti-gfp piRNA* and *cdk-1::gfp; anti-gfp piRNA; rde-3* worms. Tubulin is used as loading control.

**(E)** Schematic showing level of 22G-RNAs targeting gfp in *cdk-1::gfp*, *cdk-1::gfp; anti-gfp piRNA* and *cdk-1::gfp; anti-gfp piRNA; rde-3* worms.

**(F)** *anti-gfp piRNA*-induced 22G-RNAs were loaded into WAGOs.

**(G and H)** similar analysis to Figure 4B and C, graphs showing the fraction of GFP-positive worms in the presence of *anti-gfp piRNA* with full match at each position at the F2, F3, F4 and F5 generations (G) and with single-nucleotide mismatches at each position at the F3, F5, F7 and F9 generations (H).

**(I)** Diagram showing silent mutations of target gfp to create single nucleotide mismatches with *anti-gfp piRNA* at position t3, t15 and t21 (marked by green)

**(J)** Silent mutations of gfp appear to remain *cdk-1::gfp* expression level relative to WT.

**(K)** Western blot showing the level of CDK-1::GFP in the worms with silent mutations of gfp (left) or *anti-gfp piRNA:gfp* re-match (right).

**(L)** Nucleotides (green) showing piRNA re-match.

**(M)** Fluorescence micrographs showing the silence of CDK-1::GFP expression by piRNA re-match, see also Figure S4K (right).

**Figure S5. Related to Figure 5**.

**(A)** Percentage of dead egg and dumpy animals in WT, *prg-1 (tm872)*, and *21ur-4863* deletion strains after *sex-1* RNAi treatment.

**(B and D)** Distribution of chimeric reads (red) identified from P-CLASH and distribution of 22G-RNAs (blue) at target *fbxb-97* (B) and *comt-3* (D) in *prg-1 (tm872)* and WT. The sequence and base pairing of *21ur-1563::fbxb-97* chimera are shown (B, right) and *21ur-8264::comt-3* chimera are shown (D, right), and their locations on both of genes are indicated by the black Inverted triangles.

**(C)** RT-qPCR analysis of *fbxb-97* mRNA level in WT, *prg-1 (tm872)*, and *21ur-1563* deletion worms. actin mRNA served as the internal control. Data were collected from three independent biological replicates. Error bars represent standard deviation.

**(E)** Diagram of silent mutation of *comt-3* at four different piRNAs targeting sites shown at Figure S4D.

**(F)** Bar graphs showing the change of *comt-3* mRNA level in WT, *prg-1 (tm872)*, *comt-3::flag*, *comt-3::flag;prg-1 (tm872)*, *comt-3(*△*1p)::flag* and *comt-3(*△*4p)::flag* animals. △1p represents silent mutations of target *comt-3* to create mismatches with *21ur-8264*. △4p represents silent mutations of target *comt-3* to create mismatches with *21ur-8264*, *21ur-1363*, *21ur-2198* and *21ur-4147*. As shown in Figure S5E.

**(G)** Western blot analysis of COMT-3::FLAG level in in WT, *comt-3::flag*, *comt-3::flag;prg-1 (tm872)* and *comt-3(*△*4p)::flag* animals. Tubulin were used as loading control.

**Figure S6. Analysis of CLASH dataset and mRNA-seq, Related to Figure 6.**

**(A)** Western blot analysis of the change of protein CSR-1 in the strain (WM509) without (-) or with (+) auxin treatment. alpha-tubulin was used as loading control.

**(B)** Localization of GFP::TEV::FLAG::PRG-1 (GTF::PRG-1) in the strain (WM509) without (-) or with (+) auxin treatment.

**(C-E)** Comparison of each CSR-1 target from entire CLASH datasets. Similar analysis to Figure 6 B. Number of CSR-1 targets with piRNA targeting sites increase/decrease by 2-fold are labelled.

**(F-H)** Comparison of each CSR-1 target from RNA seq datasets. Number of CSR-1 targets increase/decrease by 2-fold are labelled.

**Table S1. *C. elegans* strains in this study (Related to Figures 1–6, Figure S1-S6)**

